# Molecular investigation of *Rlm3* from rapeseed as a broad-spectrum resistance gene against fungal pathogens producing structurally conserved effectors

**DOI:** 10.1101/2025.02.03.635517

**Authors:** Nacera Talbi, Simine Pakzad, Françoise Blaise, Bénédicte Ollivier, Thierry Rouxel, Marie-Hélène Balesdent, Karine Blondeau, Noureddine Lazar, Herman van Tilbeurgh, Carl H. Mesarich, Isabelle Fudal

## Abstract

Recognition of a pathogen avirulence (AVR) effector protein by its cognate plant resistance (R) protein triggers immune responses that render the plant resistant, representing an efficient disease control strategy. While (AVR) effectors have long been considered species- or genotype-specific, based on a lack of homology in sequence databases, a growing number of studies have now shown that these proteins belong to a limited set of structural families. This is an important finding because it paves the way for the identification or engineering of broad-spectrum R proteins capable of recognizing several members of the same structural family. In the *Leptosphaeria maculans* / rapeseed (*Brassica napus*) pathosystem, 13 *AVR* genes have been cloned, of which four encode effectors belonging to the LARS (*Leptosphaeria* AviRulence and Supressing) structural family that has also been found in thirteen other phytopathogenic fungi. Homologues of the *L. maculans* AvrLm3 AVR protein, a LARS family member, have been identified in other fungal species, including an AVR protein from *Fulvia fulva* called Ecp11-1. We have previously shown that Ecp11-1 is recognized by rapeseed varieties carrying the *Rlm3 R* gene, and that this recognition is masked by the presence of another LARS *AVR* gene, *AvrLm4-7*. In this study, we expanded our characterization of the Rlm3 resistance spectrum to putative effectors from *Fusarium oxysporum* and *Zymoseptoria ardabiliae* and showed that one effector from *F. oxysporum* f. sp. *narcissi* behaves like Ecp11-1, being recognized by Rlm3, and this recognition also being masked by the presence of *AvrLm4-7*. We finally investigated which protein regions and amino acids were necessary for the recognition of AvrLm3 and Ecp11-1 by Rlm3. This analysis is a first step towards the identification or engineering of broad- spectrum R proteins that confer protection against multiple phytopathogens through the recognition of structural effector families.

## Introduction

Breeding cultivars carrying major resistance (*R*) genes against pathogens is a powerful tool to control plant diseases (McDonald and Linde, 2002). During infection, plants carrying major *R* genes can specifically recognize pathogen effectors, which are secreted molecules that modulate plant immunity and facilitate infection (Oliva et al., 2010; Rocafort et al., 2020). Recognition of effectors, then called avirulence (AVR) proteins, triggers a set of immune responses called Effector-Triggered Immunity (ETI; (Jones and Dangl, 2006)), most often leading to a hypersensitive response (HR) characterized by a rapid cell death at the point of pathogen infection, stopping colonization of the plant. In addition to their effectiveness in protecting against pathogens, *R* genes are subject to simple genetic control, making them easy to deploy in varieties, particularly with the help of marker-assisted selection (MAS). However, the extensive deployment of single *R* genes in the field may exert strong selection pressure on the pathogens they control that become virulent through the evolution of their *AVR* gene repertoires (McDonald and Stukenbrock, 2016).

Many cases of resistance breakdown soon after the deployment of new *R* genes have been observed in the field, for example when using *R* genes against fungi responsible for cereal rusts (Kolmer, 1996; McIntosh and Brown, 1997), cereal powdery mildews (Wolfe and McDermott, 1994) or rapeseed stem canker (Rouxel et al., 2003). Different mechanisms have been reported allowing pathogens to overcome *R* genes including the acquisition of new effectors that suppress ETI, *AVR* gene deletion, inactivation, or down-regulation and point mutations that allow the virulence function of the AVR protein to be maintained while escaping recognition (Guttman et al., 2014; Jones and Dangl, 2006; Sánchez-Vallet et al., 2018). The sustainable management of *R* genes therefore represents a major challenge if we are to develop genetic control as an efficient means for limiting disease and avoiding the rapid emergence of virulent pathogen strains. Several strategies can be employed to optimize the management of *R* genes and limit the speed with which they are overcome, including the use of *R* gene combinations or alternations in the field, and combinations of major *R* genes with quantitative resistance (Brun et al., 2010). Although effective, these strategies remain difficult to implement and coordinate in the field. Another promising strategy is the use of *R* genes corresponding to effectors that are highly conserved among pathogens (called ‘core effectors’). Indeed, the durability of an R protein depends on the AVR protein it targets and on its involvement in pathogenicity. R proteins targeting AVR proteins that are strongly involved in pathogenesis would potentially be more difficult to break down, as the switch to virulence would be associated with a high fitness cost to the pathogen (Depotter and Doehlemann, 2020). In addition to their durability, *R* genes corresponding to core effectors could confer broad-spectrum resistance (BSR), as they could protect against more than one pathogen species or most races or strains of the same species (Kou and Wang, 2010). The availability of an increasing number of fungal genome sequences and effector repertoires, as well as the resolution of many fungal effector 3D structures, has led to the identification of homologous proteins and structural analogues among fungal effectors. This has, in turn, enabled the identification of *R* genes that confer resistance to a wide range of pathogens. This is notably the case for the tomato R protein Cf4, which is able to recognize the *Fulvia fulva* (formerly called *Cladosporium fulvum*) effector AVR4 but also its orthologue in *Pseudocercospora fijiensis,* or for the R protein Cf2, which confers resistance to both *F. fulva* and the nematode *Globodera rostochiensis* via their respective AVR proteins, AVR2 and Gr-VAP1. In the latter case, the effectors from the two evolutionarily distinct microorganisms target the same tomato protease guarded by Cf2 (Lozano-Torres et al., 2012; Rooney et al., 2005; Stergiopoulos et al., 2010). Structurally related effectors can also be recognized by the same resistance protein. For instance, two effectors of the MAX family, AVR-Pia and AVR1-CO39 from *Pyricularia oryzae*, are both recognized by the rice RGA4/RGA5 pair (Cesari et al., 2013; de Guillen et al., 2015).

The Dothideomycete *Leptosphaeria maculans* is responsible for stem canker (blackleg) of Brassica, notably rapeseed (*Brassica napus*). Genetic control is the main strategy to fight *L. maculans* and, as such, identification of sustainable resistance sources within Brassica species is essential. Twenty-four *R* genes against *L. maculans* (mostly called *Rlm* genes) have been genetically identified (Cantila et al., 2020; Degrave et al., 2021; Jiquel et al., 2021) and five of them have been cloned: *LepR3* (Larkan et al., 2013) and *Rlm2* (Larkan et al., 2015), which are allelic variants encoding Receptor-Like Proteins, and *Rlm9*, *Rlm4* and *Rm7* (Haddadi et al., 2021; Larkan et al., 2020), which are three allelic variants encoding Wall-Associated Kinases.

*A. L. maculans* is one of the plant-pathogenic fungi for which the largest number of AVR proteins have been identified (Balesdent et al., 2013; Degrave et al., 2021; Fudal et al., 2007; Ghanbarnia et al., 2015; Ghanbarnia et al., 2018; Gout et al., 2006; Jiquel et al., 2021; Neik et al., 2022; Parlange et al., 2009; Petit-Houdenot et al., 2019; Plissonneau et al., 2016; Van de Wouw et al., 2014), with several of these having protein homologues or structural analogues in other plant- pathogenic fungi. For example, AvrLm6 shares homologies or structural analogies with several AvrLm6-like effectors from *Venturia* species, *V. inaequalis* and *V. pirina* (Rocafort et al., 2022; Shiller et al., 2015). AvrLm10A and AvrLm10B are conserved in thirty-one plant-pathogenic fungal species from the Dothideomycetes and Sordariomycetes classes (Petit-Houdenot et al., 2019; Talbi et al., 2023). Finally, four AVR effectors of *L. maculans* (AvrLm3, AvrLm4-7, AvrLm5-9 and AvrLmS-Lep2) belong to the LARS structural family, first identified in *L. maculans*, but also present in at least thirteen other plant-pathogenic fungi (Lazar et al., 2022). Among them, AvrLm3 is homologous in sequence and structure to an AVR protein from *F. fulva*, Ecp11-1 (Mesarich et al., 2018) and both can be recognized by the same resistance protein, Rlm3, in rapeseed (Lazar et al., 2022). These data provide us with a real opportunity to investigate the possibility of developing broad-spectrum resistances against a wide range of pathogens.

*AvrLm3* is present in all *L. maculans* isolates collected in the French field, thus suggesting a central role for this effector in fungal pathogenicity (Plissonneau et al., 2017). The main mechanism allowing *L. maculans* to escape recognition by Rlm3 is the suppression of AvrLm3 recognition by another effector, AvrLm4-7, without direct physical interaction between the two AVR proteins (Plissonneau et al., 2016). This epistatic interaction was highlighted after the massive use of *Rlm7* resistance in France, and the emergence of virulent isolates that present for part of them drastic changes in *AvrLm4-*7 such as accumulation of inactivating mutations or deletion, thus abolishing the masking of *AvrLm3.* More recently, the increased use of cultivars carrying both *Rlm3* and *Rlm7* led to the selection of isolates virulent towards both genes (Balesdent et al., 2022; Plissonneau et al., 2017), with *AvrLm3* always being present in these isolates with a limited sequence polymorphism.

An AvrLm3 homologue was identified in *F. fulva*, encoded by an *AVR* gene called *Ecp11-1* that is recognized by wild tomato accessions carrying the *R* gene *CfECP11-1*. The AvrLm3 and Ecp11-1 protein sequences share 37% amino acid identity and 59% amino acid similarity (Mesarich et al., 2018), with only minor insertions/deletions between them. Lazar et al. (2022) showed that Ecp11-1 and AvrLm3 were structural analogues with five conserved disulfide bridges. Surprisingly, Ecp11-1 was able to trigger *Rlm3-*mediated resistance when introduced into a virulent *L. maculans* isolate. Furthermore, recognition by Rlm3 was masked by the presence of *AvrLm4-7*. These data suggest that *Rlm3* could be a broad-spectrum resistance source.

The present work aimed to further characterize the interaction of Rlm3 with AvrLm3 and Ecp11-1 and to determine whether Rlm3 can recognize effectors from other plant-pathogenic fungi. For this purpose, we analyzed *AvrLm3* polymorphisms in field isolates of *L. maculans* according to their phenotypic behavior towards *Rlm3*. We then further tested amino acids of AvrLm3 and Ecp11-1 potentially involved in the interaction with Rlm3 by site-directed mutagenesis. We also identified AvrLm3 homologues in proteomes of other plant-pathogenic fungi, tested whether these homologues could be recognized by Rlm3, and finally, whether this recognition could be masked by the presence of *AvrLm4-*7.

## Results

### Two amino acid changes in AvrLm3 are sufficient to escape recognition by Rlm3 in *B. napus*

We took advantage of two large *L. maculans* population surveys, which were carried out across worldwide populations of *L. maculans* by Plissonneau et al. (2017) and Balesdent et al. (2022), to decipher the interaction between AvrLm3 and its cognate R protein Rlm3. From a collection of 238 *L. maculans* isolates these surveys revealed a total of 28 polymorphic nucleotides in the coding sequence of *AvrLm3*, defining 17 different alleles that corresponded to 22 non-synonymous mutations and 17 isoforms of the protein termed AvrLm3-A to AvrLm3-R (Figure 1A). Among the polymorphic amino acid residues, three correlated quite well with the virulent and avirulent phenotypes towards *Rlm3*: I/L^58^H, G^131^R and F^134^Y (Figure 1A and B). The G^131^R change was found in all virulent alleles, while the I/L^58^H and F^134^Y changes were found in all except one virulent allele (AvrLm3-Q and AvrLm3-H, respectively). Two of these polymorphic amino acids are located on an external loop of the AvrLm3 protein (G^131^R and F^134^Y) and one is on the larger α-helix (I/L^58^H; Figure 1B).

**Figure 1.**
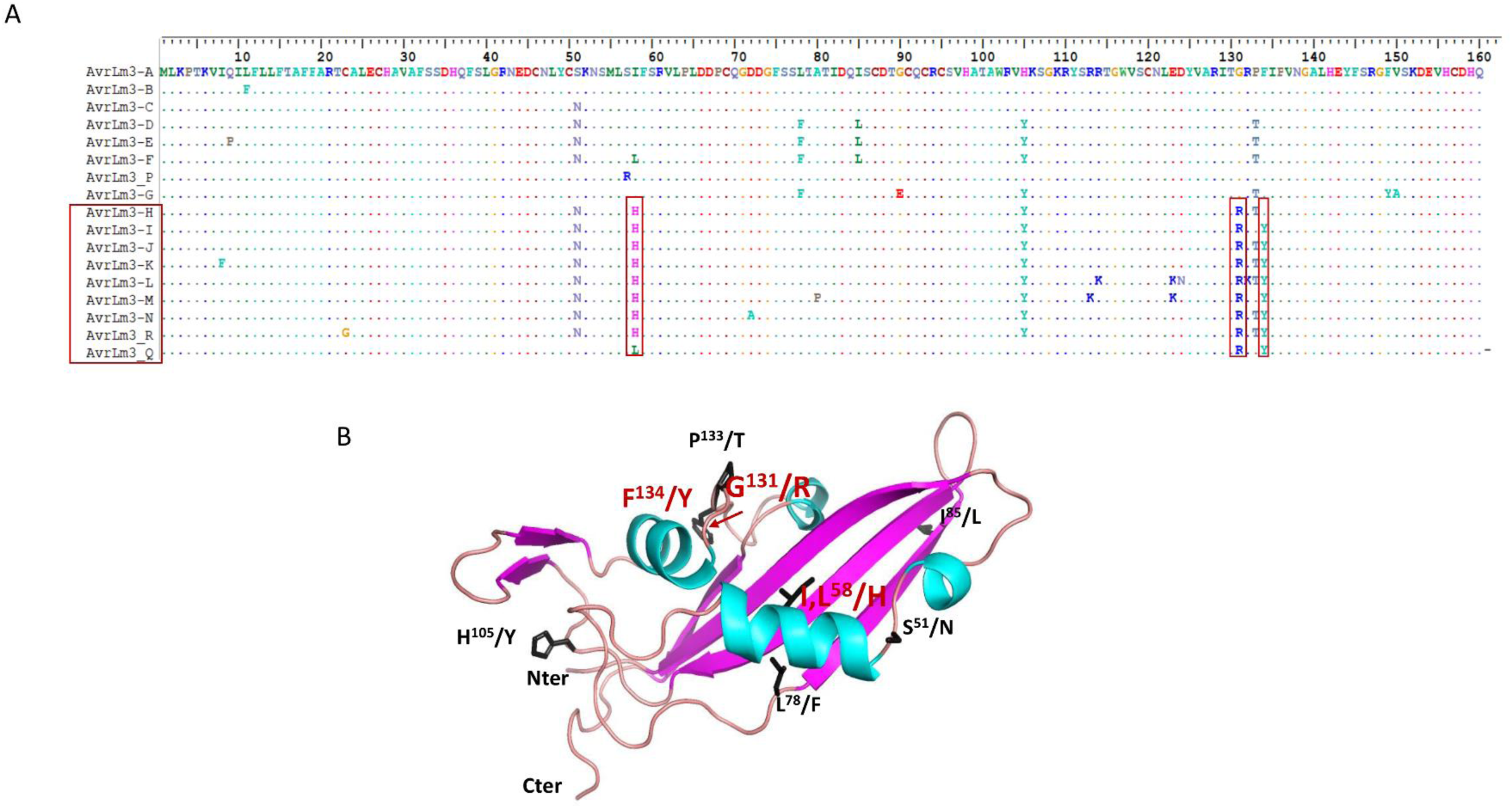
AvrLm3 alleles currently described in *L. maculans* populations and localization of polymorphic residues on the predicted 3D structure of AvrLm3. A. Amino acid polymorphisms in AvrLm3 from natural populations of *L. maculans*. Virulent alleles and conserved polymorphic amino acids identified in isolates virulent towards *Rlm3* are indicated with red boxes. Alignment was performed using BioEdit. Amino acid residues are colored according to their classes (hydrophobic - green / yellow, hydrophilic polar uncharged - orange, hydrophilic acidic - blue, hydrophilic basic – pink). B. Localisation of polymorphic residues on the 3D structure of AvrLm3 identified in populations of *L. maculans* ‘*brassicae*’. Only the polymorphic residues present in at least two alleles are shown. Amino acid changes shared among isolates virulent toward *Rlm3* are indicated in red.

To determine the role of these three residues in the induction of *Rlm3*-mediated resistance, we performed site-directed mutagenesis to generate two single mutants (AvrLm3_I58H and AvrLm3_G131R), two double mutants (AvrLm3_I58H_G131R and AvrLm3_G131R_F134Y) and one triple mutant (AvrLm3_I58H_G131R_F134Y) of AvrLm3. As these mutations involved surface-exposed residues, they are not expected to affect the global structure of AvrLm3 (Figure 2A). We generated plasmid constructs encoding the different mutated versions of AvrLm3 under the control of the *AvrLm4-7* promoter and the *ECP11-1* terminator in the binary vector pPZPnat1. The different constructs were introduced through *Agrobacterium tumefaciens*-mediated transformation into Nz-T4, an *L. maculans* isolate virulent towards *Rlm3*, *Rlm4,* and *Rlm7* (subsequently mentioned as a3a4a7, ‘a’ meaning virulent and ‘A’ avirulent). We obtained six independent Nz-T4-AvrLm3_I58H, Nz-T4-AvrLm3_G131R, and Nz-T4- AvrLm3_I58H_G131R transformants, as well as and five independent Nz-T4- AvrLm3_G131R_F134Y and Nz-T4-AvrLm3_I58H_G131R_F134Y transformants. The transformants were inoculated onto *B. napus* cv Pixel (*Rlm4*) and line 15.22.4.1 (*Rlm3*). All transformants, along with the wild-type isolate Nz-T4, were virulent on Pixel. All transformants expressing AvrLm3 with the single mutations I^58^H or G^131^R, or AvrLm3 with the double mutation I^58^H and G^131^R, were avirulent on the *Rlm3* cultivar, indicating that the amino acid mutations I^58^H and G^131^R are not sufficient to escape Rlm3-mediated recognition (Figure 3A).

**Figure 2.**
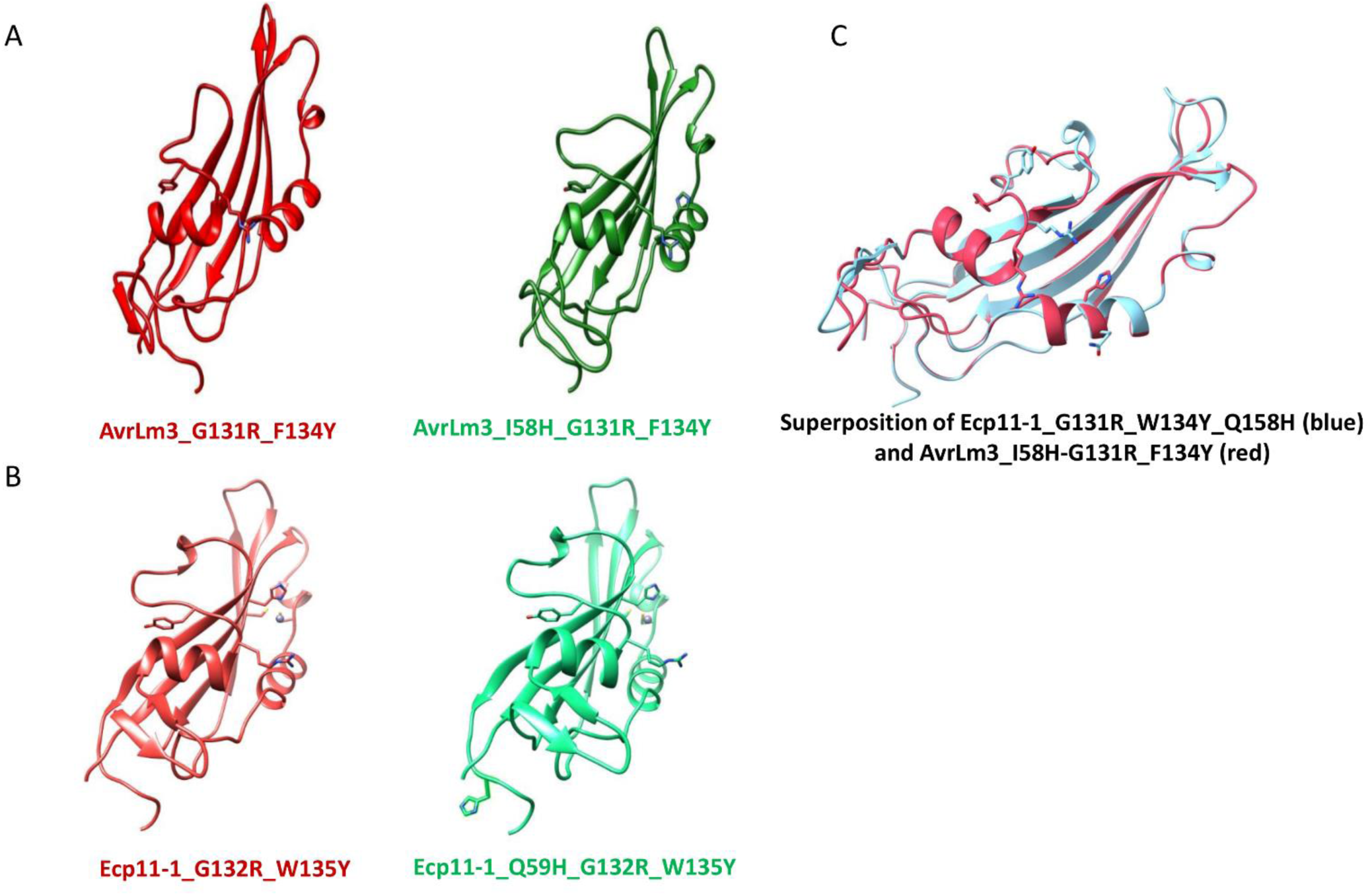
Predicted protein structures of AvrLm3 and Ecp11-1 mutated at different amino acid residues. **A.** Protein structure of mutated alleles of AvrLm3 (double mutation AvrLm3_G131R_F134Y and triple mutation AvrLm3_I58H_G131R_F134Y). **B.** Protein structure of mutated alleles of Ecp11-1 (double mutation Ecp11-1_G132R_W135Y and triple mutation Ecp11- 1_Q59H_G132R_W135Y). **C.** Superposition of the protein structures of the mutated alleles AvrLm3_I58H_G131R_F134Y and Ecp11-1_Q59H_G132R_W135Y

**Figure 3.**
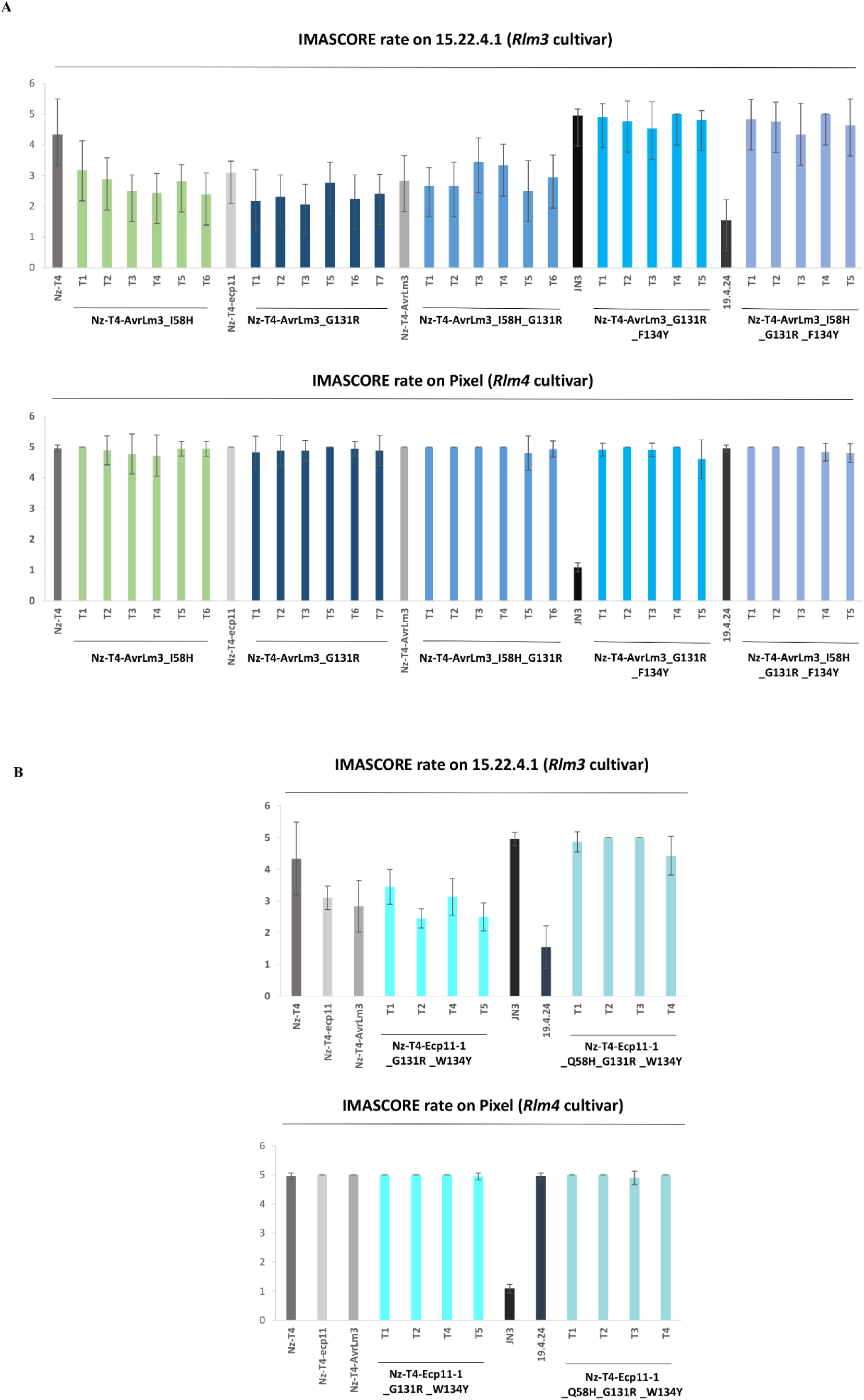
Effect of site-directed mutagenesis on AvrLm3 and Ecp11-1 recognition by Rlm3 in *B. napus*. A. Nz-T4 transformants of *L. maculans* producing AvrLm3 mutated at different amino acid residues (single mutants AvrLm3_I58H or AvrLm3_G131R, double mutants AvrLm3_I58H_G131R or AvrLm3_G131R_F134Y, and triple mutant AvrLm3_I58H_G131R_F134Y) and B. Nz-T4 transformants producing Ecp11-1 mutated at different amino acid residues (double mutant Ecp11-1_G132R_W135Y and triple mutant Ecp11-1_Q59H_G132R_W135Y) were inoculated onto cotyledons of a *B. napus* line/cultivar carrying *Rlm3* (15.22.4.1) or *Rlm4* (Pixel).

In contrast, all transformants expressing AvrLm3 with the double mutation G^131^R and F^134^Y, or with the triple mutation I^58^H, G^131^R, and F^134^Y, were virulent on the *Rlm3* cultivar. Taken together, these results suggest that the I58H mutation is not involved in the loss of recognition by *Rlm3,* but that the combined changes at residues G^131^ and F^134^ in AvrLm3 are sufficient to escape Rlm3-mediated recognition. To confirm these results, we needed to ensure that the mutated versions of *AvrLm3* leading to virulence towards *Rlm3* were expressed in isolate Nz- T4. However, since Nz-T4 (a3a4a7) naturally possesses a virulent allele of the *AvrLm3* gene, and because this gene is expressed *in planta*, a simple qRT-PCR experiment could not be performed. Thus, to confirm the expression of the mutated versions of *AvrLm3* in transformants that were virulent on the *Rlm3* line (Nz-T4-AvrLm3_G131R_F134Y and Nz-T4- AvrLm3_I58H_G131R_F134Y), a High Resolution Melting (HRM) analysis was performed. Here, the expression levels of the *AvrLm3* variants from two Nz-T4-AvrLm3_G131R_F134Y and two Nz-T4-AvrLm3_I58H_G131R_F134Y transformants were tested using cDNA derived from the susceptible *B. napus* cultivar Yudal infected with *L. maculans* at 7 days post-inoculation. Except for one transformant of Nz-T4-AvrLm3_I58H_G131R_F134Y, both the wild-type *AvrLm3* and mutated *AvrLm3* genes were expressed in all transformants tested, as revealed by the intermediate melting curves obtained when compared to the wild type isolates JN2 and Nz-T4 (Figure S1B and C).

### Three amino acid changes in Ecp11-1 are necessary to escape recognition by Rlm3 in *B. napus*

Ecp11-1 shares 37% amino acid identity with AvrLm3, with only minor insertions/deletions between the two sequences. The polymorphic residues identified in AvrLm3 that were postulated to be involved in Rlm3*-*mediated recognition correspond to changes I^58^H (Q^59^ in Ecp11-1), G^131^R (G^132^ in Ecp11-1) and F^134^Y (W^135^ in Ecp11-1) (Figure 4A). Q^59^ is located in the middle of the first α-helix and G^132^/W^135^ are in a loop that connects the second helix to the last strand (Figure 4B). To assess the involvement of Ecp11-1 residues Q^59^, G^132^ and W^135^ in recognition by Rlm3, we performed site-directed mutagenesis and generated a double mutant (Ecp11-1_G132R_W135Y) and a triple mutant (Ecp11-1_Q59H_G132R_W135Y). These mutations concern surface-exposed residues and are therefore not expected to affect the global structure of Ecp11-1 (Figure 2B). As for *AvrLm3*, constructs containing the different *ECP11-1* alleles under the control of the *AvrLm4-7* promoter and *ECP11-1* terminator in the pPZPnat1 binary vector were introduced via *A. tumefaciens*-mediated transformation into isolate Nz-T4.

**Figure 4.**
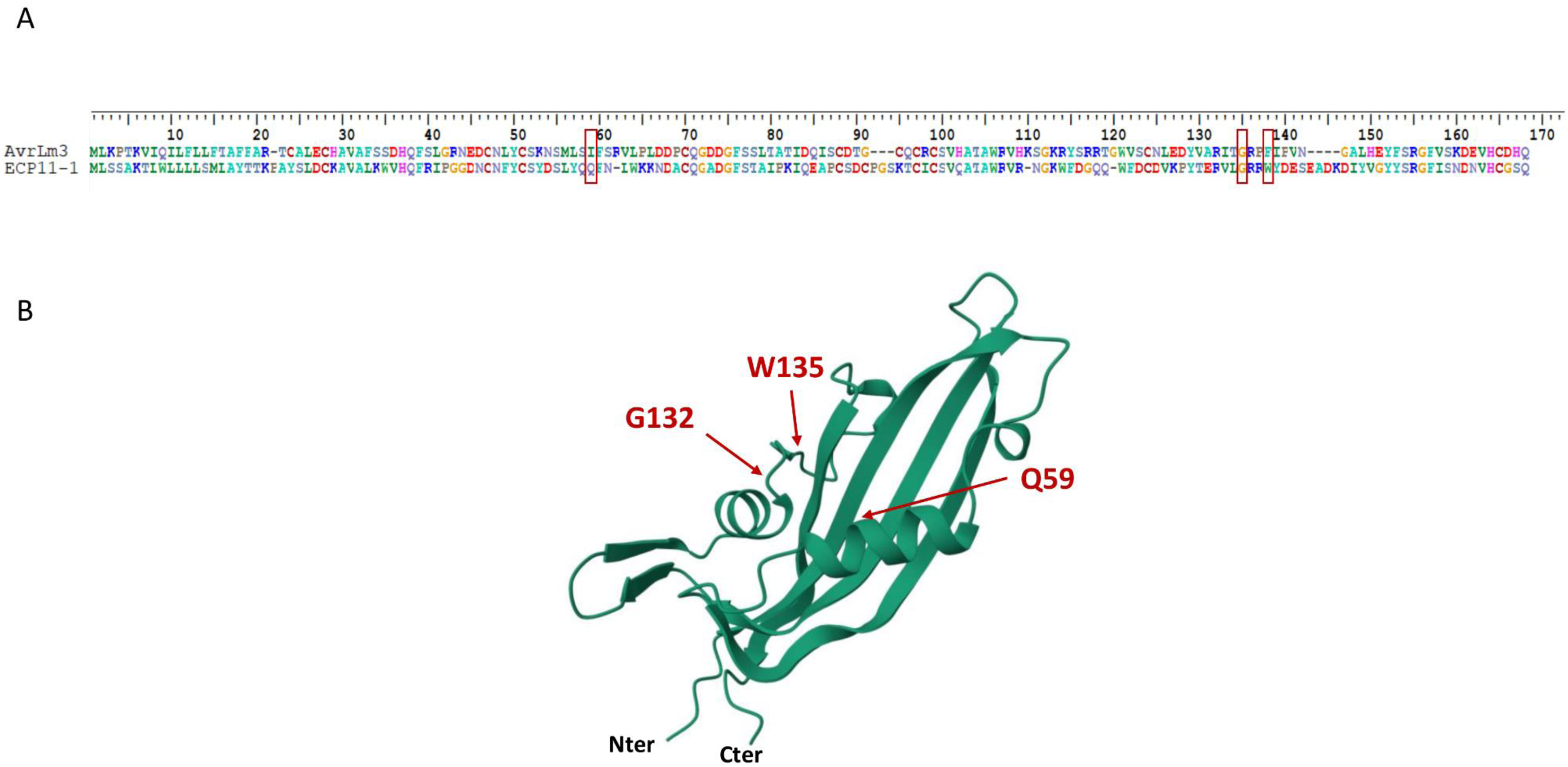
Alignment of AvrLm3 and Ecp11-1 protein sequences and localization of polymorphic residues on the Ecp11-1 3D structure. A. Alignment of AvrLm3 and Ecp11-1 protein sequences using BioEdit software. Amino acid residues are colored according to their classes (hydrophobic - green / yellow, hydrophilic polar uncharged - orange, hydrophilic acidic - blue, hydrophilic basic – pink). B. 3D structure of Ecp11-1. The residues highlighted in red correspond to the amino acids that were mutated in our study.

Wild type *L. maculans* isolates 19.4.24 (A3a4a7, ‘Ax’ meaning avirulent and ‘ax’ virulent toward the *Rlm* gene x), Nz-T4 (a3a4a7) and JN3 (a3A4A7) as well as Nz-T4 transformants carrying AvrLm3 or Ecp11-1 were used as controls. Pathogenicity was measured 12 days post- inoculation. Results are expressed as a mean score using the IMASCORE rating comprising six infection classes (IC), where IC1 to IC3 correspond to resistance, and IC4 to IC6 to susceptibility (Balesdent et al., 2005). Error bars indicate the standard deviation of technical replicates.

Four independent Nz-T4-Ecp11-1_G132R_W135Y and four Nz-T4-Ecp11- 1_Q59H_G132R_W135Y transformants were obtained. The transformants were inoculated onto *B. napus* cv Pixel (*Rlm4*) and line 15.22.4.1 (*Rlm3*). All transformants, like wild-type isolate Nz-T4, were virulent on Pixel. The transformants expressing Ecp11-1 with the double mutation Ecp11-1_G132R_W135Y were avirulent on the *Rlm3* line, indicating that the mutations G^132^R and W^135^Y are not sufficient to escape recognition by Rlm3 (Figure 3B). In contrast, all three transformants expressing the Ecp11-1 triple mutant (Ecp11- 1_Q59H_G132R_W135Y) were virulent on the *Rlm3* line. For two out of the three transformants, the expression of *ECP11-1* was validated by qRT-PCR, confirming that the gene is expressed and the protein is unable to be recognized by Rlm3 (Figure S1A). Taken together, these results suggest that the cumulated changes at residues Q^59^, G^132^ and F^135^ in Ecp11-1 are necessary to escape recognition by Rlm3.

### Homologues of AvrLm3 and Ecp11-1, with conserved 3D-structures, are present in two other phytopathogenic fungal species

As previously mentioned, Ecp11-1 is a homologue of AvrLm3 sharing 37% amino acid identity and is able to trigger *Rlm3*-mediated resistance in rapeseed. This encouraged us to search for other AvrLm3 homologues capable of triggering *Rlm3*-mediated resistance. We performed a PSI-BLAST (Position-Specific Iterative Basic Local Alignment Search Tool) search, with five iterations, against the National Center for Biotechnology Information (NCBI) nr database and identified, in addition to Ecp11-1, two more homologues, one from *Fusarium oxysporum* f. sp. *cepae* and one from *F. oxysporum* f. sp. *narcissi*. We also performed a tBLASTn search against fungal genomes available in the Joint Genome Institute (JGI) Mycocosm database and in Ensembl Fungi and identified additional homologues in *Zymoseptoria ardabiliae* and *Setosphaeria turcica.* The homologues identified in *S. turcica* corresponded to pseudo-genes and were not kept for further analysis. Taken together, we were able to find four AvrLm3/Ecp11-1 homologues in Dothideomycetes and Sordariomycetes species (Table 1). Two of them were found in the Sordariomycetes *F. oxysporum* f. sp. *narcissi* and *F. oxysporum* f. sp. *cepae* respectively, which cause basal rot of *Narcissus* and onion. The two other homologues were found in the Dothideomycete *Z. ardabiliae,* a species closely related to the wheat pathogen *Zymoseptoria tritici,* isolated from two distinct grass species: *Lolium perenne* (Ray grass) and *Dactylis glomerata* (Orchard grass). The size of three of the homologues, named Zymoseptoria_ardabiliae_STIR04_1, Zymoseptoria_ardabiliae_STIR04_2, and Fusarium_oxysporum_narcissi_BFJ63_g18418 ranges from 160 to 167 amino acids, while the fourth, Fusarium_oxysporum_cepae_BFJ70_g16777 has a size of 145 amino acids. All are enriched in cysteine residues (10 cysteines in the mature proteins) and are predicted to possess a signal peptide for secretion (Table 1).

**Table 1.** Characteristics of AvrLm3 and Ecp11-1 protein homologues identified in Fusarium oxysporum *and* Zymoseptoria ardabiliae.

| Protein name | Size (aa) | Cysteine number<sup>a</sup> | Identity with AvrLm3 | Identity with Ecp11-1 | Signal peptide<sup>b</sup> |
| --- | --- | --- | --- | --- | --- |
| AvrLm3 | 160 | 10 | - | 36% | yes |
| Ecp11-1 | 165 | 10 | 36% | - | yes |
| *Fusarium oxysporum*_narcissi_BFJ63_g18418 | 163 | 10 | 46% | 40% | yes |
| *Fusarium oxysporum*_cepaе_BFJ70_g16777 | 145 | 10 | 24% | 29% | yes |
| *Zymoseptoria*_ardabiliae_STIR04_1 | 160 | 10 | 33% | 27 % | yes |
| *Zymoseptoria*_ardabiliae_STIR04_2 | 167 | 10 | 35% | 32% | yes |

<sup>a</sup> Cysteine number is calculated based on the mature protein, without a signal peptide.
<sup>b</sup> Prediction using SignalP 3.0 software (Bendtsen et al., 2004).

The homologues shared between 24% and 46% amino acid sequence identity with AvrLm3 and between 27% and 40% with Ecp11-1 (Table 2). The highest percentage of identity (48%) was found for the two proteins from *Z. ardabiliae* (Table 2).

**Table 2.** Percentage of amino acid sequence identity (similarity) between AvrLm3, Ecp11-1 and their homologous proteins identified in Fusarium oxysporum *and* Zymoseptoria ardabiliae.

|  | Ecp11-1 | Fusarium_oxysporum_<br>narcissi_BFJ63_g18418 | Fusarium_oxysporum_<br>cepae_BFJ70_g16777 | Zymoseptoria_<br>ardabiliae_STIR04_1 | Zymoseptoria_<br>ardabiliae_STIR04_2 | 299 |
| --- | --- | --- | --- | --- | --- | --- |
|  |  |  |  |  |  | 300 |
|  |  |  |  |  |  | 301 |
| **AvrLm3** | 36% (53%) | 46% (60%) | 24% (42%) | 33% (46%) | 35% (49%) | 302 |
|  |  |  |  |  |  | 303 |
| **Ecp11-1** |  | 40% (57%) | 29% (41%) | 27% (43%) | 32% (48%) | 304 |
|  |  |  |  |  |  | 305 |
| **Fusarium_oxysporum_**<br>**narcissi_BFJ63_g18418** |  |  | 26% (40%) | 35% (51%) | 41% (54%) | 306 |
|  |  |  |  |  |  | 307 |
| **Fusarium_oxysporum_**<br>**cepae_BFJ70_g16777** |  |  |  | 30% (39%) | 26% (39%) | 308 |
|  |  |  |  |  |  | 309 |
| **Zymoseptoria_**<br>**ardabiliae_STIR04_1** |  |  |  |  | 48% (65%) | 310 |
|  |  |  |  |  |  | 311 |
|  |  |  |  |  |  | 312 |
|  |  |  |  |  |  | 313 |
|  |  |  |  |  |  | 314 |
|  |  |  |  |  |  | 315 |

The amino acid sequence identity (similarity) matrix was obtained by performing a reciprocal BLASTp analysis using the BioEdit sequence alignment editor (matrix: BLOSUM62, eValue = 1).

To get a deeper insight into the structure-function relationship of the AvrLm3/Ecp11-1 homologues, we used ColabFold, the freely accessible AlphaFold3 server. We started by generating a model for AvrLm3 which has 36% amino acid sequence identity with Ecp11-1. AlphaFold3 generated a model for AvrLm3 that was close to the Ecp11-1 structure (Figure 5). The AvrLm3 model has a four-stranded β-sheet fold covered on one side by two helical connections. This fold is characteristic of the LARS family and was observed in the crystal structures of AvrLm4-7, AvrLm5-9 and Ecp11-1. The AlphaFold3 model of AvrLm3 was also very similar to the template-based AvrLm3 model obtained from the Ecp11-1 crystal structure (Swiss-model server; Lazar et al., 2022).

**Figure 5.**
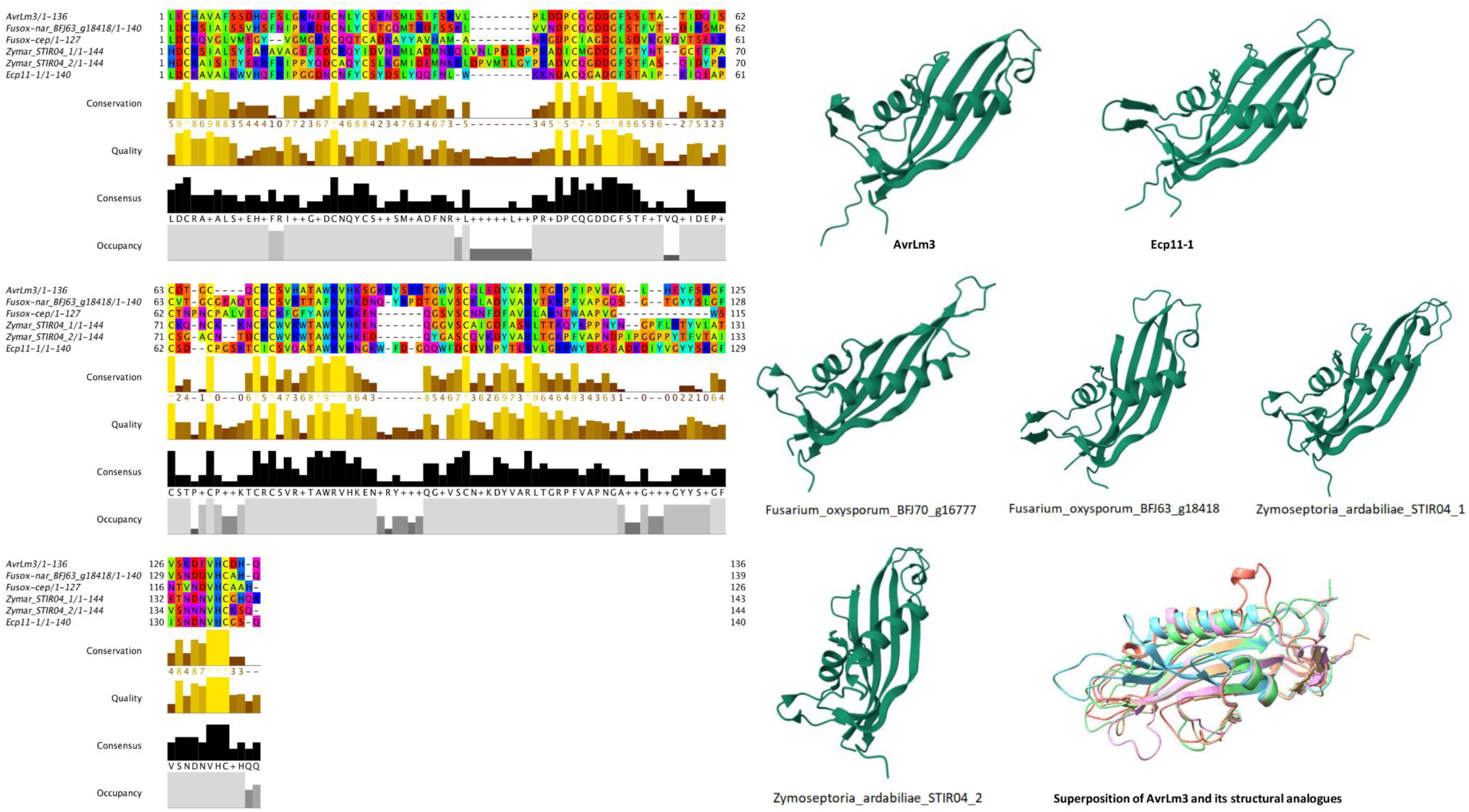
Multiple sequence alignment and predicted 3D structures of the AvrLm3/Ecp11-1 proteins and their homologues. The amino acid sequence alignment was displayed using the software Jalview (Waterhouse et al., 2009). The 3D structure predictions were performed using AlphaFold3. Ribbon diagrams of the models are presented in the same orientation.

AlphaFold3 models generated for Zymoseptoria_ardabiliae_STIR04_1, Zymoseptoria_ardabiliae_STIR04_2, Fusarium_oxysporum_BFJ63_g18418, and Fusarium_oxysporum_BFJ70_g16777 all displayed the mentioned LARS fold (Figure 5). The root mean square deviation (rmsd) values for the structural superposition of all these models with each other and with the Ecp11-1 crystal structure are presented in Table S2. The rmsd values vary between 0.7 and 0.9 Å, confirming that the overall structures of these effectors are very similar. All homologues possess ten cysteines that form superposable disulfide bridges with those from Ecp11-1. One of the ten cysteines in Zymoseptoria_ardabiliae_STIR04_1 does not align with the cysteines from the other homologues in the amino acid sequence alignment (Figure 5) but forms a disulfide bridge that superposes on those of the others (data not shown). We can therefore safely conclude that all homologues are members of the LARS family with very similar 3D structures. Divergences in the structure between the homologues are however observed in the regions that connect the β-strands.

### The AvrLm3 homologue identified in *Fusarium oxysporum* f. sp. *narcissi* triggers *Rlm3-*mediated resistance in *B. napus*

We generated constructs containing *Zymoseptoria_ardabiliae_STIR04_1*, *Zymoseptoria_ardabiliae_STIR04_2*, *Fusarium_oxysporum_BFJ63_g18418*, and *Fusarium_oxysporum_BFJ70_g16777* under the control of the *AvrLm4-7* promoter and *ECP11-1* terminator and introduced these into *L. maculans* isolate Nz-T4 via *A. tumefaciens*- mediated transformation.

We obtained four independent transformants for Nz-T4- Fusarium_oxysporum_narcissi_BFJ63_g18418, five independent transformants for both Nz- T4-Zymoseptoria_ardabiliae_STIR04_1 and Zymoseptoria_ardabiliae_STIR04_2, and six independent transformants for Nz-T4-Fusarium_oxysporum_BFJ70_g16777. The transformants were inoculated onto *B. napus* cv Pixel (*Rlm4*) and line 15.22.4.1 (*Rlm3*). All transformants, as well as wild type Nz-T4, were virulent on Pixel (Figures 6A and B). When tested on the *Rlm3* genotype, the Nz-T4-Fusarium_oxysporum_narcissi_BFJ63_g18418 transformants were avirulent, while all other transformants remained virulent (Figures 6A and B). Gene expression of all homologues was validated by qRT-PCR for one or two of the transformants, confirming that the virulence phenotype is not due to the absence or a low expression of the different genes (Figure S1A). We conclude that Fusarium_oxysporum_narcissi_BFJ63_g18418, as AvrLm3 and Ecp11-1, is recognized by Rlm3.

**Figure 6.**
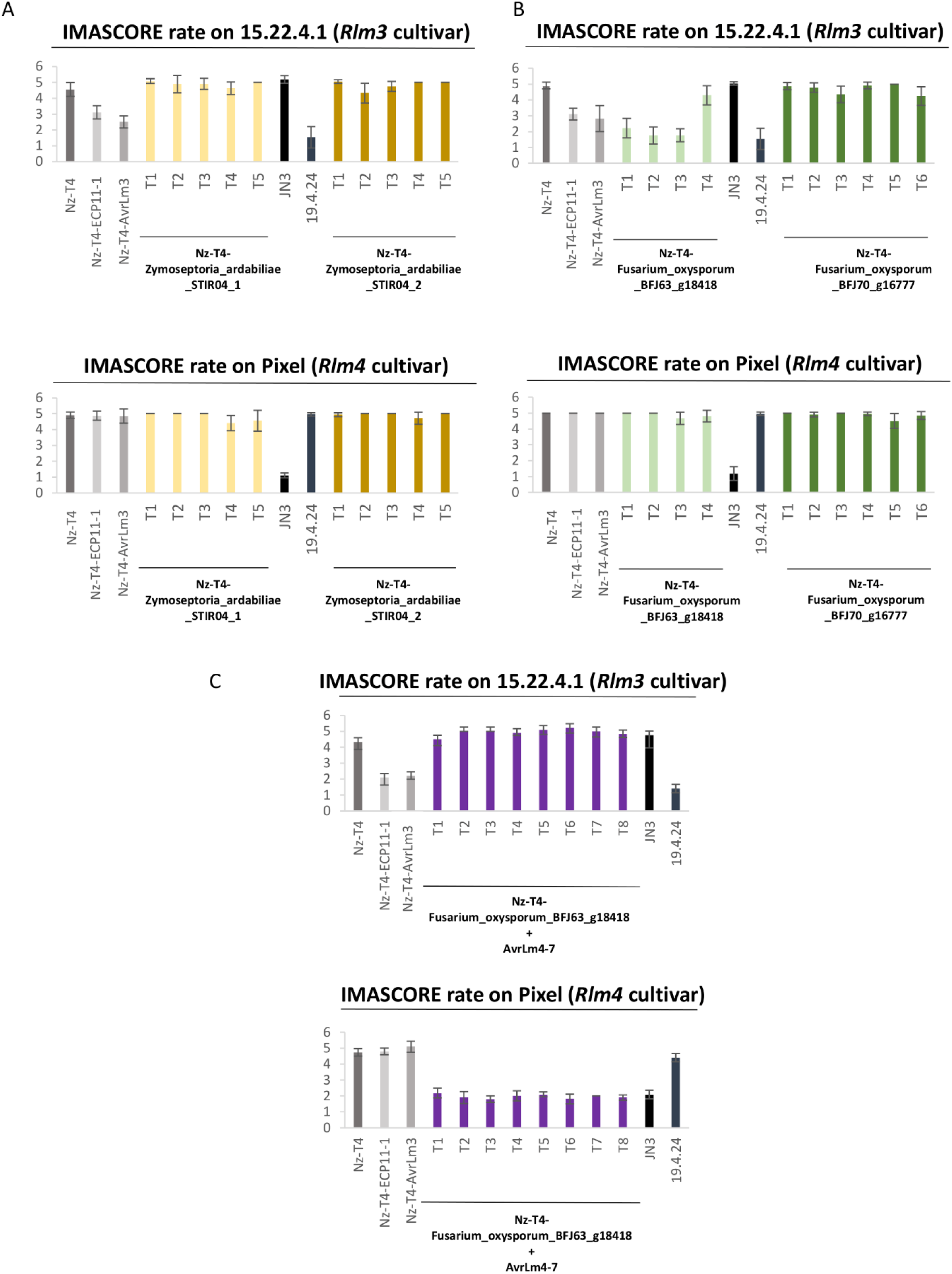
Fusarium_oxysporum_narcissi_BFJ63_g18418 triggers *Rlm3*-mediated resistance to *L. maculans* in *B. napus* that is masked by the presence of *AvrLm4-7*. Nz-T4 transformants of *L. maculans* producing A. Zymoseptoria_ardabiliae_STIR04_1 or Zymoseptoria_ardabiliae_STIR04_2 or B. Fusarium_oxysporum_BFJ63_g18418 or Fusarium_oxysporum_BFJ70_g16777 under the control of the *AvrLm4-7* promoter were inoculated onto cotyledons of a *B. napus* line/cultivar carrying *Rlm3* (15.22.4.1) or *Rlm4* (Pixel). C. Nz-T4 transformants carrying both Fusarium_oxysporum_BFJ63_g18418 and *AvrLm4-7* were inoculated onto cotyledons of a *B. napus* cultivar carrying *Rlm3* (15.22.4.1) or *Rlm4* (Pixel). For all inoculation tests, wild type of *L. maculans* isolates 19.4.24 (A3a4a7, ‘Ax’ meaning avirulent and ‘ax’ virulent toward the *Rlm* gene x), Nz-T4 (a3a4a7) and JN3 (a3A4A7) as well as Nz-T4 transformants producing AvrLm3 or Ecp11-1 were used as controls. Pathogenicity was measured 12 days post-inoculation. Results are expressed as a mean score using the IMASCORE rating comprising six infection classes (IC), where IC1 to IC3 correspond to resistance, and IC4 to IC6 to susceptibility (Balesdent et al., 2005). Error bars indicate the standard deviation of technical replicates.

### The *Rlm3-*mediated resistance triggered by the AvrLm3 homologue from *Fusarium oxysporum* f. sp. *narcissi* is masked by the presence of *AvrLm4-7*

Lazar et al. (2022) previously showed that the presence of *AvrLm4-7* suppressed the ability of Ecp11-1 to trigger *Rlm3*-mediated resistance. We assessed whether the presence of *AvrLm4-7* was also able to suppress the ability of Fusarium_oxysporum_BFJ63_g18418 to induce *Rlm3-* mediated resistance. Thus, we complemented a Nz-T4- Fusarium_oxysporum_BFJ63_g18418 transformant (transformant T3) with *AvrLm4-7* using the plasmid pBht2-AvrLm4-7 previously constructed by Lazar et al. (2020). We obtained eight independent Nz-T4- Fusarium_oxysporum_BFJ63_g18418 transformants for which the presence of *AvrLm4-7* was confirmed by PCR (data not shown). We inoculated the transformants onto the *B. napus* cultivars Pixel (*Rlm4*) and 15.22.4.1 (*Rlm3*), as well as onto an *Rlm7* cultivar (Figure 6C and data not shown). While wild type Nz-T4 was virulent on Pixel (*Rlm4*) and *Rlm7* cultivars, the transformants were avirulent on both cultivars (Figure 6C and data not shown), confirming that *AvrLm4-7* was expressed in the transformants. While the transformants expressing Fusarium_oxysporum_BFJ63_g18418 were avirulent on the *Rlm3* cultivar, the Nz-T4- Fusarium_oxysporum_BFJ63_g18418-AvrLm4-7 transformants were virulent (Figure 6B-C). We conclude that the presence of *AvrLm4-7* suppresses recognition of Fusarium_oxysporum_BFJ63_g18418 by Rlm3.

## Discussion

In this study, we investigated the ability of the Rlm3 R protein from rapeseed to recognize AvrLm3 from *L. maculans* and its homologues from other plant-pathogenic fungi. Using site-directed mutagenesis, we identified amino acids involved in the recognition of AvrLm3 and Ecp11-1 by Rlm3. We further demonstrated the ability of Rlm3 to recognize a homologue of AvrLm3 found in *F. oxysporum* f. sp. *narcissi*, as well as the ability of AvrLm4-7 to suppress that recognition. These data strongly suggest that Rlm3, which targets AvrLm3, a ‘core’ effector conserved in isolates of *L. maculans* and several other plant-pathogenic fungi, could be used for broad-spectrum resistance.

Making use of *AvrLm3* polymorphism data collected from natural populations of *L. maculans*, several polymorphic residues correlating with the avirulent or virulent phenotype towards *Rlm3* were identified. Three polymorphic residues (I/L^58^H, G^131^R and F^134^Y) were found in all but two virulent alleles of AvrLm3 (AvrLm3-Q and AvrLm3-H), with these two alleles not having the I/L^58^H or F^134^Y polymorphism (but a P^133^T polymorphism), respectively. Interestingly, the projection of the polymorphic amino acid positions of AvrLm4-7 and AvrLm5-9 onto their 3D structures proved to be informative. Only three polymorphic isoforms of AvrLm5-9 were reported in *L. maculans* populations (Ghanbarnia et al., 2018), including two point mutations at residue R^55^ to either T or K leading to virulence towards *Rlm9*. Similarly, Parlange et al. (2009) identified one point mutation in AvrLm4-7 at residue G^120^ to R that is responsible for virulence towards *Rlm4.* Residue I^58^ is located in the same region of the AvrLm3 structure as residue R^55^ in AvrLm5-9. Similarly, residue G^131^ is located in the same region of the AvrLm3 structure as residue G^120^ (responsible for the switch to virulence towards *Rlm4*) in the AvrLm4- 7 structure, suggesting the same protein regions could be involved in the virulence phenotypes. These residues could be part of regions that are in contact with the corresponding R proteins. Notably, Haddadi et al. (2021) cloned *Rlm4* and *Rlm7* and found they were allelic to *Rlm9* (Larkan et al., 2020), while in another study, *Rlm3* was found to be closely linked to these *R* genes and possibly allelic (Delourme et al., 2004). This suggests that, collectively, the R proteins encoded by these genes will have almost identical 3D structures and therefore could interact similarly with their respective cognate AVR proteins. We demonstrated that the double mutation G^131^R/F^134^Y in AvrLm3 was sufficient to escape recognition by Rlm3. Consequently, residue I/L^58^ does not appear to be necessary for recognition of AvrLm3 by Rlm3, which is consistent with the polymorphism in *L. maculans* populations, since a virulent allele (Avrlm3- Q) of AvrLm3 carried the L^58^ residue. However, we found that the corresponding residue in Ecp11-1 (Q^59^) was involved in the recognition of Ecp11-1 by Rlm3. Indeed, only the triple mutant Q^59^H/G^132^R/W^135^Y allowed Ecp11-1 to escape recognition by Rlm3 and not the G^132^R/W^135^Y double mutant. The involvement of a third amino acid in the recognition of Ecp11-1 by Rlm3 could be explained by a slightly different interacting region allowing a less efficient recognition by Rlm3, whether that interaction is direct or indirect. Indeed, although the mutations are not predicted to affect the global structure of Ecp11-1, they could, nevertheless, reduce the interacting surface. For example, the tomato serine/threonine protein kinase Pto is guarded by the R protein Prf and is targeted by two AVR proteins from *Pseudomonas syringae* pv. tomato, AvrPto and AvrPtoB (Kim et al., 2002). AvrPto and AvrPtoB are not homologues, and their 3D structures are different. Nevertheless, the crystal structure of the complexes between AvrPtoB and Pto and between AvrPto and Pto revealed that both AVR proteins bind to the same protein surface of Pto but that the detailed interactions are not similar (Dong et al., 2009). In the same way, distinct alleles of the AVR protein AvrL567 from *Melampsora lini* are recognized by the R proteins L5, L6 and L7 from flax through a direct interaction. 3D structures of two alleles of AvrL567 revealed that polymorphic amino acids were located at the protein surface, changing affinity for their R proteins cumulatively (Wang et al., 2007).

We found four homologues of AvrLm3 and Ecp11-1, with no predicted function, in other plant-pathogenic fungal species: two in *Z. ardabiliae* and two in two *formae speciales* of *F. oxysporum.* Even if distantly related, *Z. ardabiliae*, *L. maculans* and *F. fulva*, belong to the Dothideomycetes class, and the presence of AvrLm3 homologues in these three species could more or less be expected. It was more of a surprise to find homologues in a much more distant species as *F. oxysporum*, which belongs to the Sordariomycetes class. Nevertheless, this is not the first time that homologous effector/AVR proteins of *L. maculans* have been found in *F. oxysporum*: AvrLm2 shows sequence homologies with Six1 (also known as Avr3; Ghanbarnia et al., 2015) and the AvrLm10A/AvrLm10B pair was found to be conserved in several *Fusarium* species (Petit-Houdenot et al., 2019; Talbi et al., 2023). It is interesting to note that all four species, *L. maculans, F. fulva*, *F. oxysporum* and *Z. ardabiliae*, share the same hemibiotrophic and apoplastic / vascular lifestyle, which may influence their effector repertoire, and the conservation of common effectors acting during asymptomatic colonization. The amino acid sequence identity levels (between 23% and 43%) are highly significant considering these effectors are under strong selection pressure to evolve rapidly. Moreover, the AlphaFold3 predictions of the 3D structure of these homologues confirm they all have the LARS fold consisting of a four-stranded β-sheet covered by helices and loops on one side. Each of the AvrLm3 homologues has 10 cysteines involved in superposable disulfide bridges. These observations suggest a common evolutionary origin.

Among the four homologous proteins expressed in *L. maculans*, only Fusarium_oxysporum_narcissi_BFJ63_g18418, was recognized by Rlm3. This is not surprising since the protein shares the highest amino acid sequence identity with AvrLm3 and Ecp11-1 (46% and 40%, respectively). However, Fusarium_oxysporum_narcissi_BFJ63_g18418 shares only one of the residues we found to be involved in the recognition by Rlm3, F^122^ (corresponding to F^134^ in AvrLm3 and to W^135^ in Ecp11-1). Focusing on the residues involved in the recognition by Rlm3, Zymoseptoria_ardabiliae_STIR04_2 appears as a good candidate since it shares two of these residues with AvrLm3 (G^131^ and F^134^). However, although all the homologues transformed into *L. maculans* were highly expressed, Zymoseptoria_ardabiliae_STIR04_2 was not able to induce *Rlm3-*mediated resistance, suggesting that conservation of the overall 3D-structure is also crucial for recognition. The structure of Zymoseptoria_ardabiliae_STIR04_2 is very similar to the crystal structure of Ecp11-1 and to the model of AvrLm3. Although all homologues possess the same fold, notable differences exist in the conformations of the connections between the strands, where many of the polymorphic positions are situated. Sequence alignment of the AvrLm3 homologues revealed some highly conserved peptides (^48^DGV, ^80^WRV and ^135^VH, Ecp11-1 numbering). Interestingly these sequences form a conserved patch on one side of the β-sheet. This patch is not near the residues that were identified to be important for the recognition by Rlm3 but could hint at a region that is involved in the interaction between the effector and its host virulence target. In the absence of structures of complexes with their target or R proteins, it remains very difficult to speculate why one homologue is recognized by Rlm3 but not the others.

In summary, this study reinforces that *Rlm3* could be used as a broad-spectrum *R* gene recognizing LARS effector family members. We have now identified three effectors from three distinct plant-pathogenic fungal species recognized by Rlm3. However, neither *F. fulva* nor *F. oxysporum* f. sp. *narcisii* are pathogenic of rapeseed, but, interestingly, genera of the Sordariomycetes (including *Fusarium*) and Dothideomycetes classes (including *Cladosporium*) have been identified in the microbiota associated with rapeseed during *L. maculans* infection (Kerdraon et al., 2019; Kerdraon et al., 2020). As soon as the corresponding genomes have been sequenced, it will be interesting to look for AvrLm3 homologues and members of the LARS structural family in these species and study their interaction with Rlm3. Homologues of AvrLm3 and LARS effectors in different plant-pathogenic fungi could also be used to identify *R* genes in the corresponding host plants, as potential new sources of resistance. Hypothesizing that *Rlm3* could potentially be allelic to *Rlm4*, *Rlm7* and *Rlm9*, and encode a WAKL protein, the R proteins recognizing AvrLm3 homologues in other host plants, such as CfEcp11-1 in wild tomatoes, could also correspond to WAKL proteins, which would help to identify these proteins among R protein candidates. Alternatively, Rlm3 could be transferred from rapeseed into a host plant of pathogens carrying *AvrLm3* homologues.

## Materials and Methods

### *Leptosphaeria maculans*, bacteria and plant growth conditions

The *L. maculans* Nz-T4 isolate (virulent towards *Rlm3*, *Rlm4* and *Rlm7*) is a field isolate from New Zealand (Parlange et al., 2009) that was used to perform all *A. tumefaciens*-mediated transformations. The *L. maculans* v23.1.3 isolate (avirulent towards *Rlm4* and *Rlm7*, and virulent towards *Rlm3*; also known as JN3) originated from *in vitro* cross #23 (Balesdent et al., 2001) and was used as a control in pathogenicity tests. Fungal cultures were maintained on V8 (vegetable juice) agar medium at 25°C for 7 days in the dark for mycelial growth and between 10 and 14 days under a mixture of white and near-UV light for sporulation (Ansan-Melayah et al., 1995). *L. maculans* conidia were harvested for transformation and incubated in Fries liquid medium (yeast extract 5 g/L, NH4 5 g/L, sucrose 30 g/L, KH2PO4 1 g/L, NH4NO3 1 g/L, MgSO4 500 mg/L, CaCl2 130 mg/L, NaCl 100 mg/L) for 24 h at 28°C under agitation (200 rpm) to enable germination.

*Escherichia coli* strain DH5α and *A. tumefaciens* strain C58 pGV2260 were grown on lysogeny broth (LB) medium (peptone 10 g/L, yeast extract 5 g/L, NaCl 10 g/L) containing the appropriate antibiotics at the appropriate concentration: rifampicin 25 µg/mL, ampicillin 50 µg/mL, kanamycin 100 µg/mL. The *E. coli* strains were incubated for 24 h at 37°C, while the

*A. tumefaciens* strains were incubated for 48 h at 28°C.

*B. napus* plants were grown in chambers set to a 16 h light (22 °C, 80% humidity):8h dark (18 °C, 100% humidity) cycle.

### Fungal transformation

*A. tumefaciens*–mediated transformation was performed on *L. maculans* as described by Gout et al. (2006). Transformants were selected on minimal medium containing nourceothricin (50mg/mL) or hygromycin (50 µg/mL) and cefotaxime (250mg/mL), and purified by isolating individual pycnidia onto the same selective medium upon sporulation.

### Inoculation tests on rapeseed

A cotyledon inoculation test, as described by Balesdent et al. (2002), was used to phenotype *L. maculans* isolates and transformants generated for their virulence towards *Rlm3*, *Rlm7* and *Rlm4* rapeseed genotypes. Spore suspensions (10^7^ pycnidiospores/ml) of each isolate or transformant were inoculated on 10–12 plants of the *B. napus* Pixel (*Rlm4*), 15.22.4.1 (*Rlm3*) and 18.22.6.1 (*Rlm7*) cultivars. Symptoms were scored 12–21 days post-inoculation (dpi) using a semi-quantitative 1 (avirulent) to 6 (virulent) rating scale, with a score of 1 to 3 representing different levels of resistance (from typical HR to delayed resistance) and 4 to 6 representing different levels of susceptibility (mainly evaluated by the intensity of sporulation on lesions (Balesdent et al., 2005). Pathogenicity tests were repeated twice.

### Plasmid constructs and DNA manipulation

The sequences of interest (*AvrLm3* and *ECP11-1* with mutations in one to three residues of interest, and the four *AvrLm3* homologues identified in *F. oxysporum* and *Z. ardabiliae*) were synthesized and cloned into a pTwist_amp vector by Twist Bioscience (San Francisco, California, USA). Restriction sites were integrated on both sides of the different genes of interest by the supplier (*Eco*RI/*Xho*I or *Sal*I/*Xho*I).

The plasmid pPZPnat1-promAvrLm4-7 was used for both the cloning of constructs and the transformation of *L. maculans*. It corresponds to the plasmid pPZPnat1 which contains a kanamycin resistance cassette for the selection of transformed bacteria (*A. tumefaciens* and *E. coli*) and a nourceothricin resistance gene used for the selection of fungal transformants (Gardiner et al., 2005) to which the *AvrLm4-7* promoter was added (Lazar et al., 2022).

Constructs allowing expression of the genes of interest under the control of the *AvrLm4-7* promoter and the *ECP11-1* terminator were generated using the GIBSON cloning kit (New- England Biolabs, Evry, France). The pPZPnat1-promAvrLm4-7 vector was digested with *Eco*RI and *Xho*I or *Sal*I and *Xho*I (in the case of Zymoseptoria_ardabiliae_STIR04_1) according to the supplier’s recommendations. In parallel, the inserts corresponding to the genes of interest and the terminator of *ECP11-1* were amplified by PCR using Phusion Taq polymerase (Invitrogen, Carlsbad, USA) under suitable PCR conditions and specific primers constructed according to the recommendations of the cloning kit (Table S1). The inserts were then integrated into the pPZPnat1-promAvrLm4-7 vector using NEBuilder GIBSON DNA assembly according to the supplier’s recommendations. The plasmid pBht2-AvrLm4-7, containing the *AvrLm4-7* gene, promoter and terminator, as well as the hygromycin resistance gene used for the selection of fungal transformants, was previously constructed by Lazar *et al*. (2022).

### RNA extraction, reverse transcription, and qRT-PCR analysis

RNA of *B. napus* cotyledons infected with *L. maculans* was extracted using TRIzol® Reagent (Invitrogen, Cergy Pontoise, France) and cDNA was generated using oligo-dT-primed reverse transcription with PrimeScript Reverse Transcriptase (Clontech, Palo Alto, CA, U.S.A.) as described by Plissonneau et al. (2016). To investigate transgene expression in transformed *L. maculans* isolates, qRT-PCR was performed using a Bio-Rad CFX96 Touch Real-Time PCR Detection System and ABsolute SYBR Green ROX dUTP Mix (ABgene, Courtaboeuf, France) as described by Fudal et al. (2007). Here, Ct values were analyzed as described by Muller et al. (2002) to evaluate transgene expression level. For each value measured, two technical replicates and two biological replicates were assessed. All primers used for qRT-PCR are detailed in Table S1.

### HRM experiments and analyses

HRM was performed to analyze expression levels in transformed isolates of *L. maculans* containing two copies of *AvrLm3* differing by a few nucleotides. The HRM mix was composed of 2 μL of cDNA template, 4 μL of each primer (2 μM, Table S1) and 10 μL of SsoFast™ EvaGreen® Supermix (Bio-Rad, Hercules, CA, United States). PCR amplification was carried out with a Bio-Rad CFX96 Touch Real-Time PCR Detection System according to the manufacturer’s instructions: an initial denaturation step at 98°C for 2 min, followed by 40 cycles at 98°C for 5 s, 60°C for 5 s and a final melting step from 60°C to 95°C with an increase of 0.2°C every 5 s. An HRM curve analysis was performed by collecting data from the melting step and results were analysed with Bio-Rad Melt Curve Analysis™ Software.

### Bioinformatics analysis and structure prediction

To search for sequence homologues of AvrLm3, a PSI-BLAST analysis was performed against the NCBI nr database. The analysis was complemented with a tBLASTn search against fungal genomes available in the JGI Mycocosm database and on Ensembl Fungi using default parameters. Signal peptide was predicted using SignalP 3.0 software (Bendtsen et al., 2004).

We generated 3D structural models for the AvrLm3 homologs with AlphaFold3 (Jumper et al., 2021; Varadi et al., 2021) using the Colab serverhttps: //alphafoldserver.com. The reliability of the structure predictions was assessed by the Local Distance Difference Test (LDDT) score, as reported by the programs. Structural models were analyzed with PyMOL v2.5.1 (Schrödinger, LLC) or Chimera (Pettersen et al., 2004). It should be mentioned that, at the time of this work, the crystal structures of Ecp11-1 and AvrLm5-9 were not in the Protein Data Bank (PDB) database.

Sequence alignment used the web service functionalities of the Jalview platform (Waterhouse et al., 2009).

## Acknowledgements

The authors wish to thank all members of the “Effectors and Pathogenesis of *L. maculans*” group; Anaïs Pitarch and Laetitia Dupont for help with plasmid cloning; the greenhouse technician staff for plant management; the administrative supporting staff for the administrative and financial follow-up of this project, and the laboratory glassware staff of the BIOGER research unit. N. Talbi was funded by a PhD salary from the University Paris-Saclay. The “Effectors and Pathogenesis of *L. maculans*” group benefits from the support of Saclay Plant Sciences-SPS (ANR-17-EUR-0007). This work was supported by the Plant Health and Environment Division of INRAE (AAP 2019 Resistrans) and the French National Research Agency project STARlep (ANR-20-CE20-0026).

## Supporting information legends

**Figure S1:**
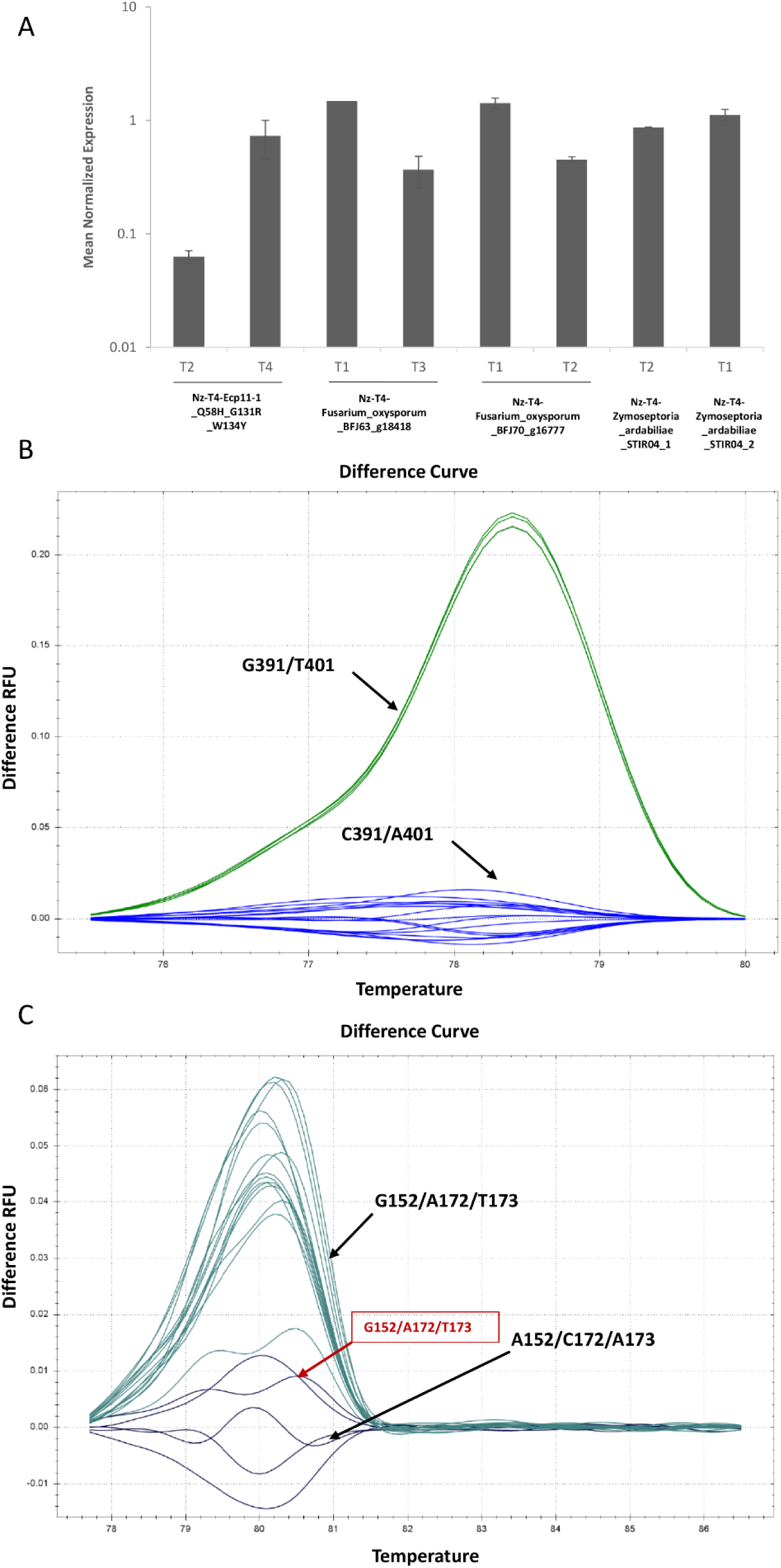
Relative expression of *AvrLm3* and its homologues in *L. maculans* transformants during infection of *B. napus*. **A.** Relative expression of *AvrLm3* homologues in Nz-T4 transformants of *L. maculans*. RNA extractions were performed on infected cotyledons of cv. Yudal at 7 dpi by *L. maculans* Nz-T4 isolate transformed with Ecp11-1_Q58H_G131R_W134Y, Fusarium_oxysporum_BFJ63_g18418, Fusarium_oxysporum_BFJ70_g16777, Zymoseptoria_ardabiliae_STIR04_1 and Zymoseptoria_ardabiliae_STIR04_2. One or two transformants were analyzed per transformation. Gene expression levels are relative to *L. maculans Actin* and calculated as proposed by Muller *et al*. (2002). Each data point is the average of two biological replicates and two technical replicates. Standard error of the mean normalized expression level is indicated by error bars. **B.** and **C**. Clustering of melting curves of *AvrLm3* amplified in wild type *L. maculans* isolates JN2 and Nz-T4 and in the Nz-T4 transformants complemented with different alleles of AvrLm3 and virulent on *Rlm3* cultivar: an AvrLm3 allele mutated for two amino acids (AvrLm3_G131R_F134) or three amino acids (AvrLm3_I58H_G131R_F134Y). RNA extraction was performed on cotyledons of cv. Yudal infected by JN2, Nz-T4 and four transformants at 7 dpi. Primers used for HRM are summarized in Table S1: Primers used for B. allowed two polymorphic nucleotides to be distinguished: G^391^C and T^401^A. Isolates displaying the *AvrLm3* G^391^/ T^401^ allele (JN2 isolate) are clustered in yellow and isolates displaying the AvrLm3 C^391^/ A^401^ allele, including Nz-T4, Nz-T4-AvrLm3_G131R_F134Y and Nz-T4- AvrLm3_I58H_G131R_F134Y are clustered in blue. Primers used for C. allowed tree polymorphic nucleotides to be distinguished: A^152^G/C^172^A/A^173^T. Isolates displaying the *AvrLm3* A^152^/C^172^/A^173^ allele, including Nz-T4 (wild type for *AvrLm3*) and one transformant AvrLm3_G131R_F134 (T2) (indicated by the grey arrow) are clustered in purple and isolates displaying the AvrLm3 G^152^/A^172^/T^173^ allele, including JN2, Nz-T4- AvrLm3_G131R_F134Y (T1) and Nz-T4-AvrLm3_ I58H_G131R_F134Y are clustered in dark green. RFU, Relative Fluorescence Unit.

**Table S1.**
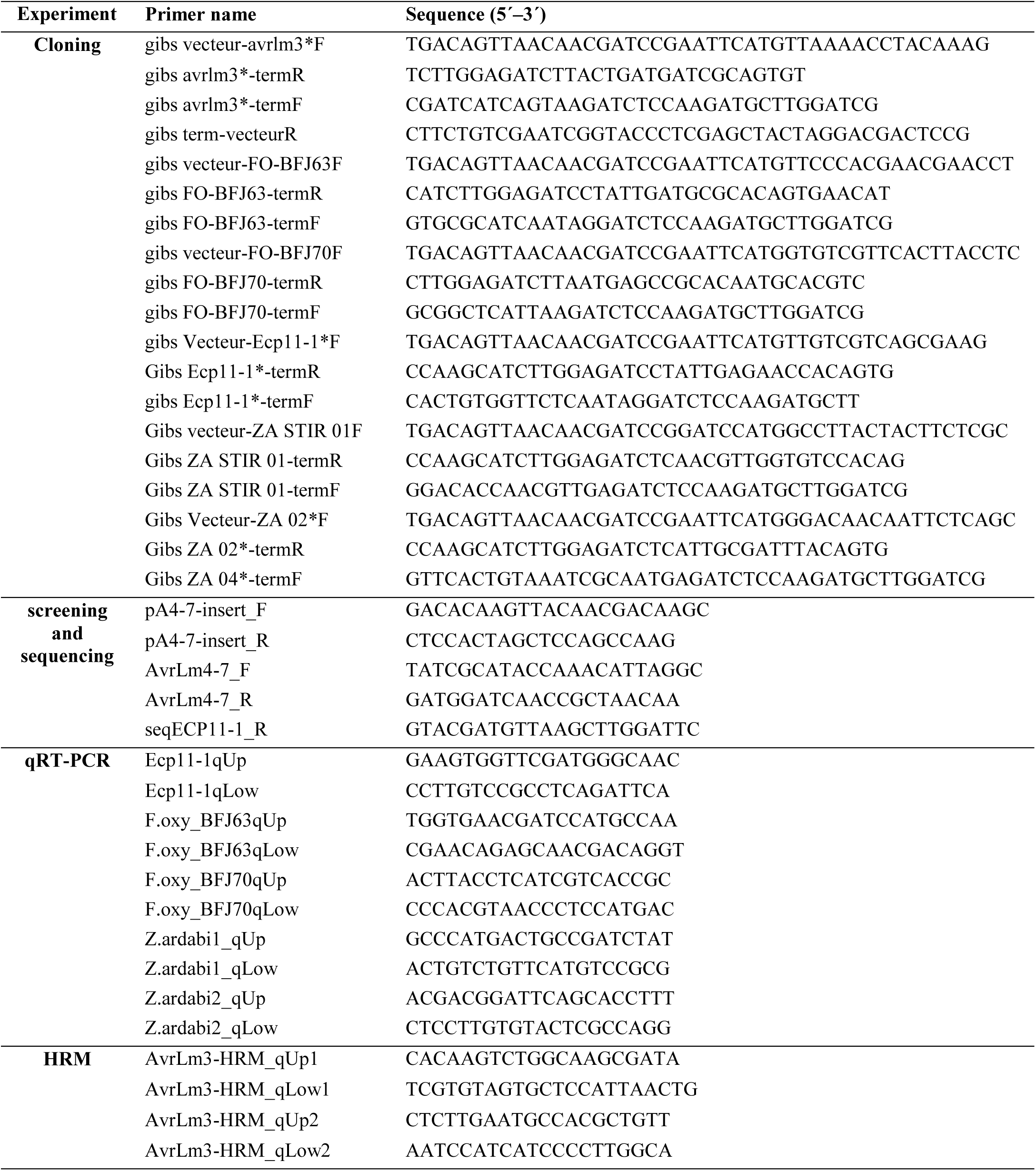
Primers used in this study.

**Table S2.** Comparison between the protein structures of AvrLm3 homologues.

|  | Ecp11 | AvrLm3 | Zar-1 | Zar-2 | Foxnar | LDDT |
| --- | --- | --- | --- | --- | --- | --- |
| AvrLm3 SM | 0.8A / 106 Ca / 37% |  |  |  |  | 90 |
| Zar-1 | 0.8A / 94 Ca / 24% | 0.7A / 90 Ca / 34% |  |  |  | 90 |
| Zar-2 | 0.7A / 111 Ca / 32% | 0.8A / 90 Ca / 34% | 0.7A / 108Ca / 45% |  |  | 90 |
| Foxnar | 0.8A / 114Ca / 37% | 0.7A / 115 Ca / 43% | 0.8A / 99Ca / 34% | 0.7A / 114 Ca / 41% |  | 90 |
| Foxcep | 0.9A / 55Ca / 23% | 0.9A / 56 Ca / 20% | 0.7A / 63 Ca / 20% | 0.8A / 59Ca / 24% | 0.9A / 60Ca / 26 % | 90 |

AvrLm3 AF 1.2A/22Ca/37% alpha fold model.
AvrLm3 SM SWISS-MODEL.
LDDT: Local Distance Difference Test model confidence score (above 90, very high confidence; between 70 and 90, high confidence; between 50 and 70, low confidence; lower than 50, very low confidence).

